# Inter- and intracellular colonization of Arabidopsis roots by endophytic actinobacteria and the impact of plant hormones on their antimicrobial activity

**DOI:** 10.1101/222844

**Authors:** Anne van der Meij, Joost Willemse, Martinus A. Schneijderberg, René Geurts, Jos M. Raaijmakers, Gilles P. van Wezel

## Abstract

Many actinobacteria live in close association with eukaryotes like fungi, insects, animals and plants. Plant-associated actinobacteria display (endo)symbiotic, saprophytic or pathogenic life styles, and can make up a substantial part of the endophytic community. Here, we characterised endophytic actinobacteria isolated from root tissue of *Arabidopsis thaliana* (Arabidopsis) plants grown in soil from a natural ecosystem. Many of these actinobacteria belong to the family of *Streptomycetaceae* with *Streptomyces olivochromogenes* and *Streptomyces clavifer* as well represented species. When seeds of Arabidopsis were inoculated with spores of *Streptomyces* strain coa1, which shows high similarity to *S. olivochromogenes*, roots were colonised intercellularly and, unexpectedly, also intracellularly. Subsequent exposure of endophytic isolates to plant hormones typically found in root and shoot tissues of Arabidopsis led to altered antibiotic production against *Escherichia coli* and *Bacillus subtilis*. Taken together, our work reveals remarkable colonization patterns of endophytic streptomycetes with specific traits that may allow a competitive advantage inside root tissue.

## INTRODUCTION

Actinobacteria represent a diverse phylum composed of both rod-shaped and filamentous bacteria that can be found in soil, marine and fresh water ecosystems (Goodfellow 2012). The filamentous actinobacteria are versatile producers of bioactive natural products, including two-thirds of all known antibiotics as well as many anticancer, antifungal and immunosuppressive agents (Barka et al. 2016; Bérdy 2005; Hopwood 2007). The actinobacteria are also major producers of industrially relevant enzymes (Anne et al. 1990; van Dissel et al. 2014). Hence, the actinobacteria are of utmost importance for human health, agriculture and biotechnology.

The concept of actinobacteria as free-living bacteria has more recently been challenged by studies pointing to their intimate relationships with diverse eukaryotes (Seipke et al. 2012; van der Meij et al. 2017). Indeed, they have been found in association with vertebrates, invertebrates, fungi and plants. Actinobacteria are welcome guests to their hosts due to their ability to produce chemically diverse natural products. Much of the chemical diversity of secondary metabolites produced by actinobacteria has likely evolved because of their interactions with other (micro)organisms in highly diverse environments (Seipke et al. 2012; van der Meij et al. 2017). It is becoming increasingly clear that actinobacteria play a key role in maintaining plant health by contributing to biotic and abiotic stress tolerance (Viaene et al. 2016). For example, actinobacteria produce siderophores for iron acquisition as well as antibacterials and antifungals to protect their host against pathogens (Viaene et al. 2016).

Actinobacteria can make up a substantial part of the root endophytic community across the plant kingdom, which is largely determined by an increased relative abundance of the family of *Streptomycetaceae* (Bulgarelli et al. 2012; Edwards et al. 2015; Lundberg et al. 2012). The composition of rhizospheric and root endophytic bacterial communities is strongly influenced by soil type (Bulgarelli et al. 2012) as well as by the plant genotype (Perez-Jaramillo et al. 2017). Therefore, part of the microbiome composition of the rhizosphere and endosphere of *Arabidopsis thaliana* grown under controlled conditions in different natural soils is conserved with specific bacterial taxa including members of the actinobacteria (Bulgarelli et al. 2012). Despite their occurrence in the endosphere, very little is known on how most actinobacteria colonise the endosphere and where they reside. An exception is *Streptomyces scabies*, the causal agent of potato scab, which has been studied in detail (Loria et al. 2006; Jourdan et al. 2016; Bignell et al. 2010; Bukhalid et al. 1998).

Colonization of root and shoot tissue by actinobacteria is dependent, at least in part, on chemical cues from the plant as was shown for enrichment of actinobacteria by the plant hormone salicylic acid (SA) (Lebeis et al. 2015). SA plays a role in a variety of physiological and biochemical processes and is important as an endogenous signal mediating local and systemic plant defense responses against pathogens and abiotic stress factors (Rivas-San Vicente and Plasencia 2011). SA is detectable in Arabidopsis leaves and roots at concentrations up to 1 μg/g fresh weight (van de Mortel et al. 2012). Hence, endophytic actinobacteria are most likely exposed and responsive to SA and other plant hormones such as jasmonic acid (Halim et al. 2006; Zhao 2010).

In this study we isolated endophytic actinobacteria from Arabidopsis roots, grown under controlled conditions as well as from plants grown in an ecological setting. Colonization patterns were monitored and visualised by confocal as well as electron microscopy for *Streptomyces* strain coa1, which shows high similarity with *Streptomyces olivochromogenes*, on axencially-grown *A. thaliana*. Finally, we investigated how specific plant hormones affect the antimicrobial activity of endophytic *Streptomyces* isolated from wild *Arabidopsis* providing a first step towards evaluating the concept of plant-mediated ‘antibiotic production on demand’.

## RESULTS & DISCUSSION

### Isolation of endophytic actinobacteria

To confirm the presence of actinobacteria in the endosphere, sterile *A. thaliana* Col-0 plants were grown in a potting soil:sand mixture for two weeks under controlled conditions, followed by harvesting roots and shoot, surface-sterilization and homogenization of the root tissue and plating onto various media selective for actinobacteria (Zhu et al. 2014b). Ten morphologically distinct actinobacterial isolates were obtained. A similar sampling approach was adopted for *A. thaliana* ecotype mossel (Msl) obtained from a natural ecosystem (Mossel, Veluwe, the Netherlands). Next to plating onto selective media, total DNA was extracted in eight replicates from samples obtained from the soil, the rhizosphere, the root endosphere and from a toothpick inserted in the soil. these DNA samples were then analyzed by 16S rDNA-amplicon sequencing. Based on total sequence reads the results of the latter analysis showed that actinobacteria represented on average 22% of the total endophytic population (Fig. 1A). Among the actinobacterial operational taxonomic units (OTUs), the *Streptomycetaceae* and *Micromonosporaceae* were overrepresented. Additionally, 10% the reads of the endophytic population was represented by only two OTUs (137 and 48), belonging to the family of *Streptomycetaceae* (Fig. 2B). This enrichment was not as much observed for the soil, the rhizosphere and bacteria associated with a toothpick, suggesting a species-specific selection among the root endophytic actinobacteria of *A. thaliana*. Numbers on total sequence reads and OTUs are listed in Table S1.

**Figure 1.**
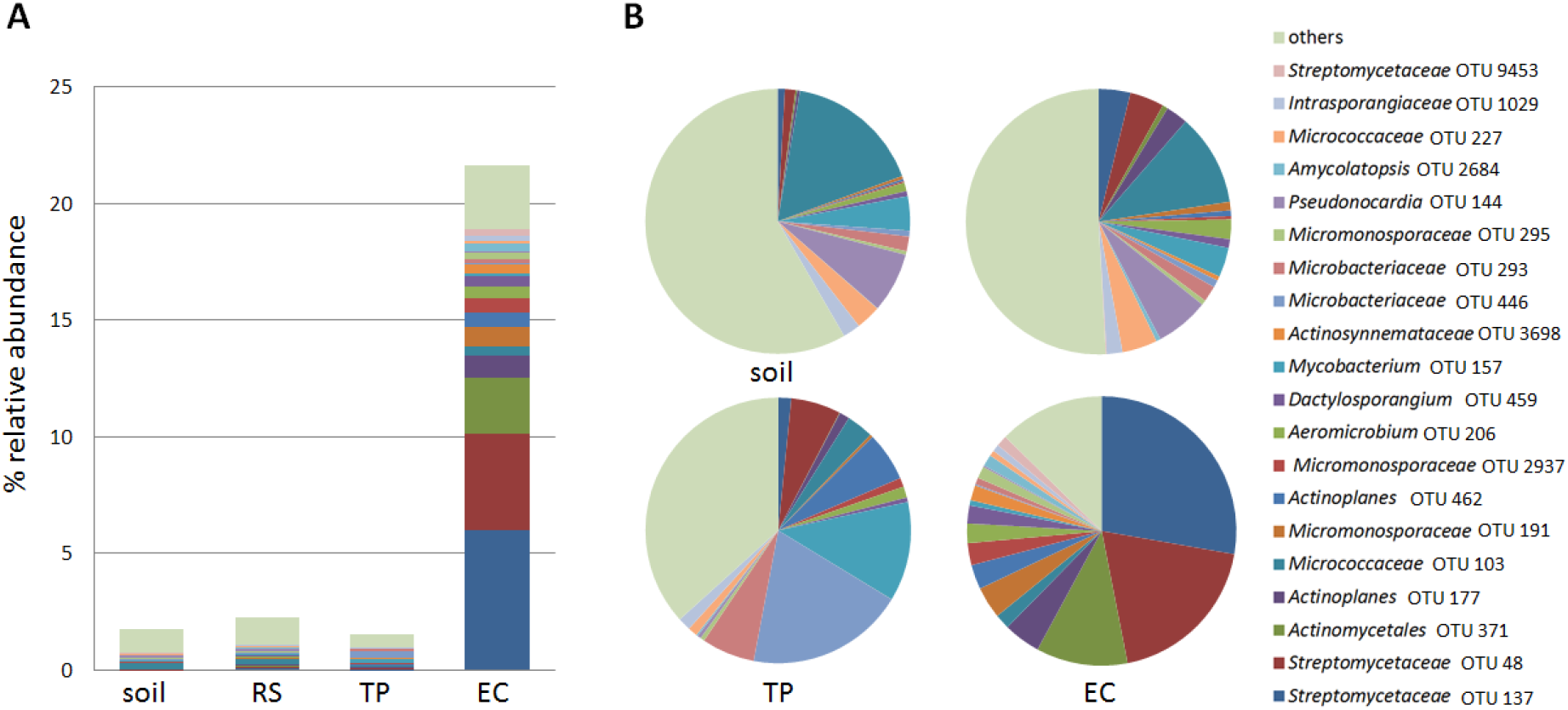
Relative abundance of the actinobacteria in the endophytic compartment of *A. thaliana* Msl. A) Amplicon sequencing data show that actinobacteria represent 22 % of the total endophytic compartment (EC), while in soil and rhizosphere (RS) communities and those in association with a toothpick (TP) inserted in the soil, they take up roughly 2 % of the total bacterial community. B) The enrichment of actinobacteria is predominately driven by *Streptomycetaceae* OTU 48 and 137. OTU 48 and 137 make up 10% of the total EC and almost 50% of all the actinobacteria present in the EC. These OTUs are not as much enriched in the samples derived from non-endophytic origins.

Isolation of actinobacteria from the endosphere *A. thaliana* ecotype Msl resulted in 35 morphologically different isolates. The 34-465 region of the 16S RNA gene of all isolates was sequenced and the closest hits on the EzBioCloud were confirmed using the MLSA-based phylogeny published recently (Labeda et al. 2017). Linking the OTUs detected by 16S rDNA amplicon sequencing to a single *Streptomyces* species was not always possible due to a relatively low resolution (Girard et al. 2013; Labeda et al. 2017). Nevertheless, the two isolates that were analysed both belonged to the most abundant OTUs *Streptomyces clavifer* and *S. olivochromogenes* (Fig. 2A). Out of the 35 isolates, five were closely related to *S. clavifer* and four to *S. olivochromogenes*. Supportive evidence comes from several independent studies, reporting *S. olivochromogenes* from the endosphere of Chinese cabbage roots, potato tubers, medicinal plants and purple henbit (Singh and Gaur 2016; Lee et al. 2008; Doumbou et al. 1998; Kim et al. 2012). Additionally, strains similar to *S. clavifer* were previously isolated from sugarcane, lucerne plants and wheat (Kruasuwan et al. 2017; Franco et al. 2017; Misk and Franco 2011).

**Figure 2.**
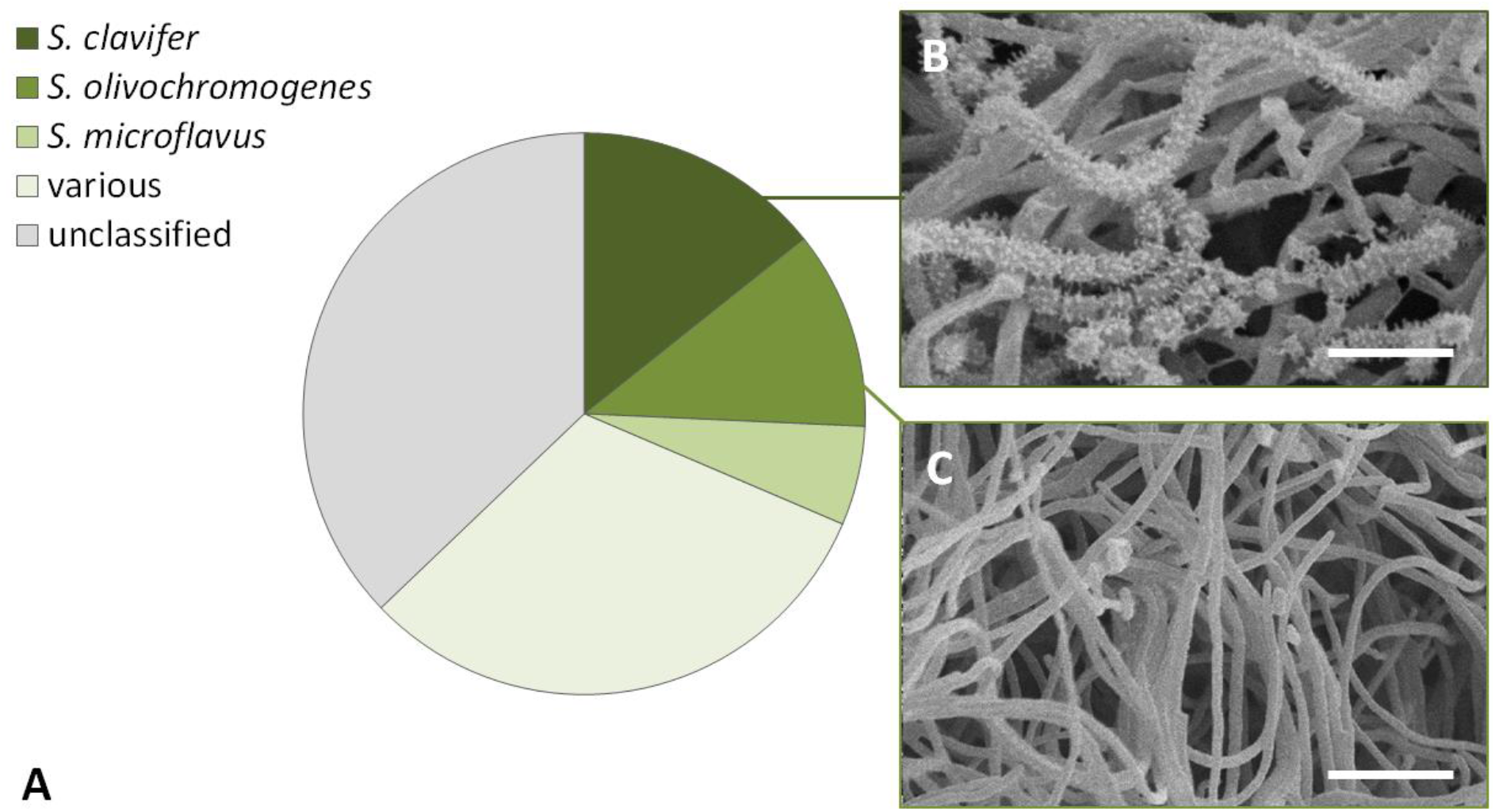
Characterization of *Streptomyces* endophytes and their taxonomic distribution. A) Several isolates show highest similarity with *Streptomyces clavifer* or *Streptomyces olivochromogenes* on the basis of 16S rDNA analysis. The majority of the isolates showed a wide variation in closest species assigned, indicating a diverse endophytic compartment. ‘Unclassified’ means that these actinobacteria could not be classified at the species level based on the 16S rDNA sequence. B) Scanning electron micrograph of *Streptomyces* sp. MOS18, which produces spiny spores. Scale bar: 3 μM. C) Scanning electron micrograph of *Streptomyces* sp. MOS38 showing poor sporulation. Scale bar: 5 μM. Strains were grown on SFM media for 6 days.

Subsequent phenotyping of these endophytic actinobacterial isolates by high resolution imaging by scanning electron microscopy (SEM) revealed spiny spores that are typical of *S. clavifer* (Goodfellow 2012), supporting the 16S rDNA-based taxonomic classification of the isolates as *S. clavifer* (Fig. 2B). In contrast, *S. olivochromogenes* poorly develops sporulating mycelium according to Bergey’s, which is shown as well for an isolate classified as *S. olivochromogenes* (Fig. 2C). However, impaired sporulation was encountered frequently, a phenotype that was dependent on the growth media, which makes this feature non-specific (see next paragraph). The endophytic isolates of *A. thaliana* ecotype Msl showed diverse colony morphologies (for examples see Fig. 3). Remarkably, 20 out of 35 isolates failed to develop on R5 agar plates, suggesting they had either lost key sporulation genes or lacked the ability to develop in the absence of specific nutrients or trace elements. For example, R5 agar is known to lack sufficient iron and copper, which is a major reason why *bld* mutants cannot sporulate on this media (Keijser et al. 2000; Lambert et al. 2014). Additionally, several endophytes showed the propensity to open up their colony, with their vegetative mycelium facing upward. This feature is well exemplified by *Streptomyces* sp. MOS31 and MOS18 (Fig. 3H and C). MOS18 is taxonomically related to *S. clavifer* and MOS31 is the only endophytic isolate closely related to *Streptomyces bobili*, a ‘neighbour’ of *S. olivochromogenes* (Labeda et al. 2017). To study the morphological characteristics of MOS31 at higher resolution SEM was applied, which revealed a thick sheet of hyphae that turned away from the inside of the colony (Fig. 4A). Additionally, we observed hyphae extending from these sheets, away from the vegetative mycelium (Fig. 4B).

**Figure 3.**
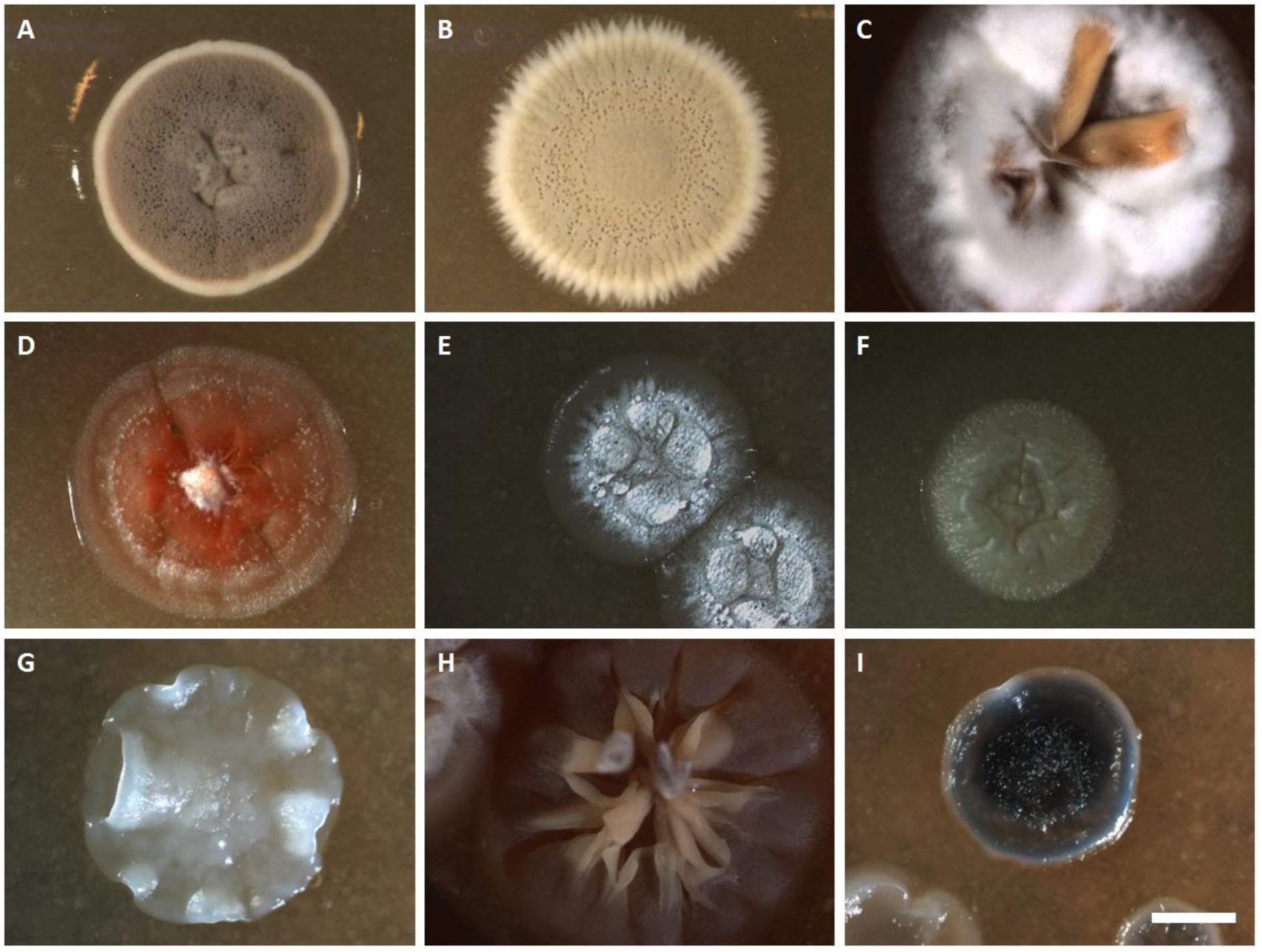
*Streptomyces* endophytes display a wide range of morphologies. Strains shown above are grown on SFM agar plates for 6 days. *S. coelicolor* M145 and *S. griseus* DSM40236 are shown as reference strains. A) *S. coelicolor* M145 B) *S. griseus* DSM40236 and the endophytic streptomycetes MOS18 (C), MOS38 (D), MOS14 (E), MOS32 (F), MOS25 (G), MOS31 (H) and MOS35 (I). Scale bar: 2 mm.

**Figure 4.**
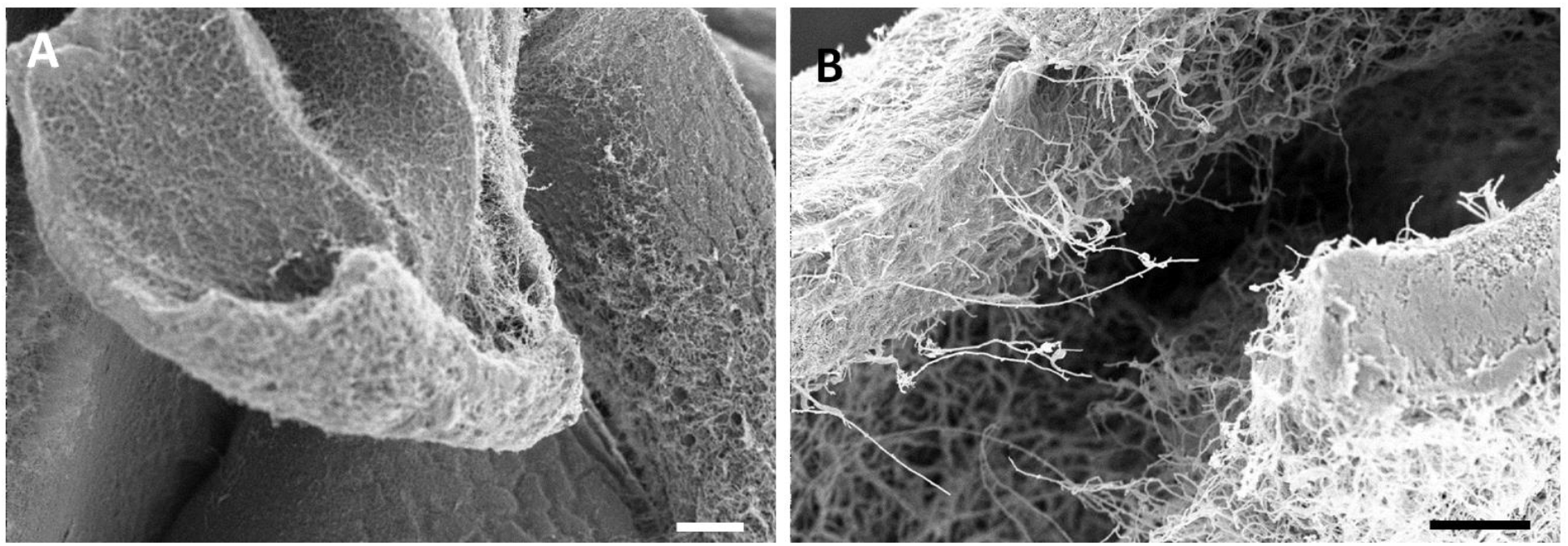
Scanning electron micrograph of *Streptomyces* sp. MOS31. A) Thick layers of vegetative mycelium made up of hyphae and extracellular matrix turn away from the inner part of the colony. Scale bar: 100 μM. B) Hyphae extending from the mycelial sheets, thereby growing away from the vegetative mycelium. Scale bar: 30 μM. *Streptomyces* sp. MOS31 was grown on SFM agar plates for 6 days.

### Endophytic colonization of *A*. thaliana by *Streptomyces*

We then wanted to know if and where the isolates enter the root endosphere of *A. thaliana* to get more insight into the yet elusive endophytic biology of *Streptomyces*. Therefore we inoculated spores of *S. olivochromogenes* strain coa1, which was recruited by sterile *A. thaliana* Col-0 plants grown in a potting soil:sand mixture, onto sterilised seeds of *A. thaliana* Col-0. Attachment of the spores to the seeds was confirmed by SEM (Fig. S1). Seven-day-old seedlings grown from the treated seeds were stained with propidium iodide and subjected to confocal fluorescence microscopy. The results showed that the endophyte had colonised both leaves and roots (Fig. 5). Strain coa1 attached to the lateral roots in dense pellets, whereas colonization of the leaves involved hyphal growth over the leaf surface. The colonised *Arabidopsis* roots were then fixed for sectioning and high resolution imaging with transmission electron microscopy (TEM). Regions of interest were identified by obtaining 1-μm sections that were stained with toluidine to visualise the bacteria (Fig. 6A). The results showed that coa1 not only colonised the root surface but also the internal root tissue and, remarkably, the intracellular space (Fig. 6BC). No plant cell defects were observed in the imaged samples. Strikingly there is no plant cellular membrane separating *Streptomyces* from the intracellular space.

**Figure 5:**
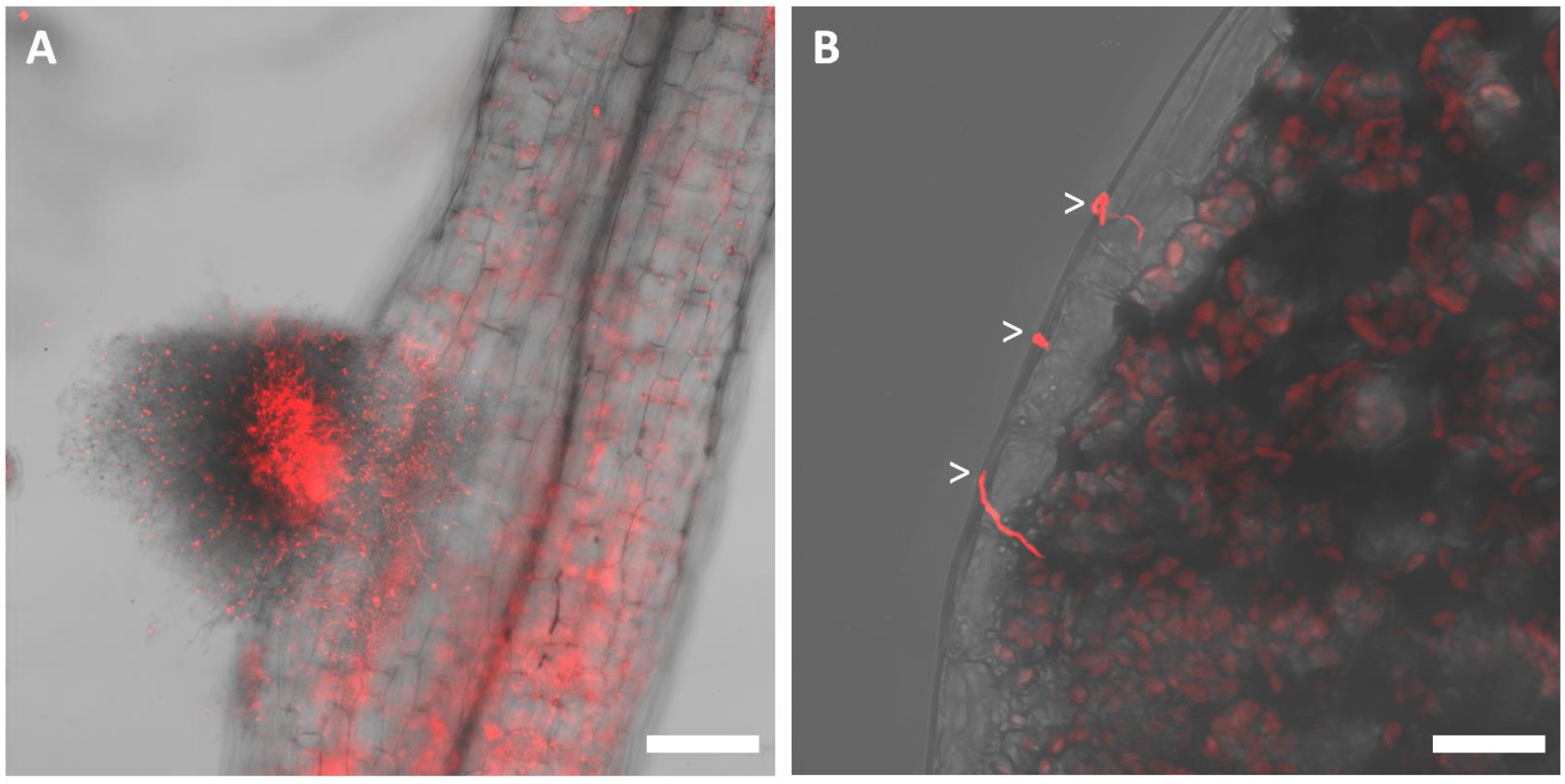
Colonization of *Arabidopsis* by *Streptomyces* endophyte coa1. A) Confocal micrograph of a colonised lateral root. The sample is stained with propidium iodide, resulting in red fluorescence of both bacterial and plant cells. Coa1 attaches to the root as a dense pellet. Scale bar: 50 μM. B) Confocal micrograph of the border of the leaf. Single hyphae are growing over the leaf surface (arrowheads). Scale bar: 15 μM.

**Figure 6:**
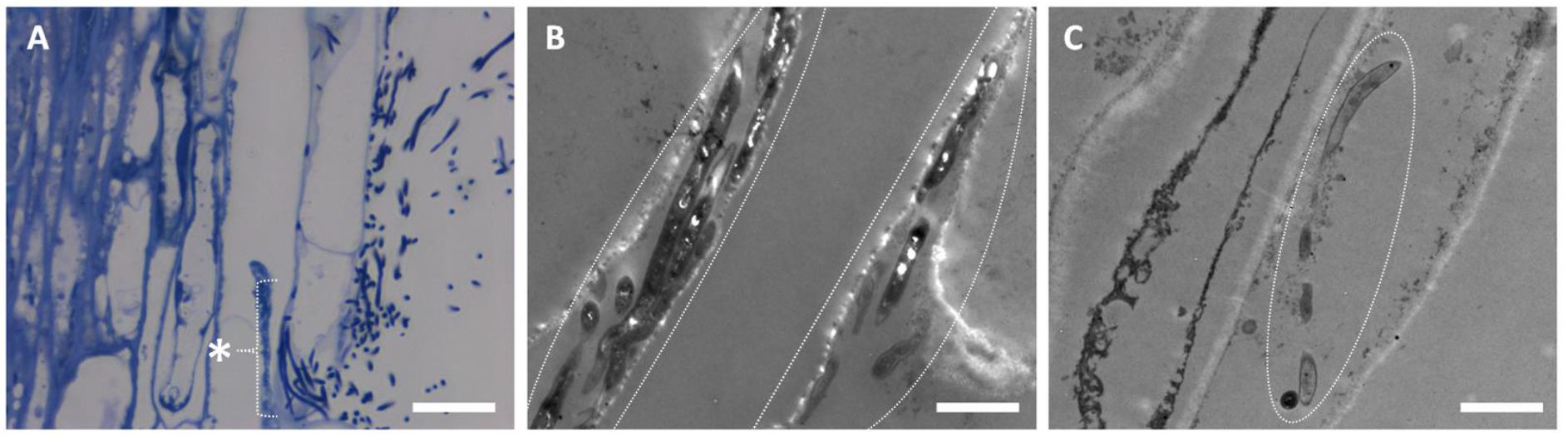
Sections of *Arabidopsis* roots invaded by *Streptomyces* coa1. A) Toluidine stained section of an *Arabidopsis* root invaded by *Streptomyces* coa1. The bacterium enters the root via intercellular space (asterisk). Scale bar: 10 μM. B & C) transmission electron micrographs of *Arabidopsis* roots. Image B shows intercellular presence of the bacterium (between dashed lines) whereas image C shows intracellular growth of *Streptomyces* coa1 in *Arabidopsis* root cells (oval). Scale bar: 2 μM.

Previous studies showed colonization of the root surface of *Arabidopsis* by actinobacteria, and lettuce, turnip rape and carrot by *Streptomyces* (Bulgarelli et al. 2012; Bonaldi et al. 2015; Kortemaa et al. 1994). In addition, endophytic colonization of germinating wheat seed has been reported by reintroduction of a GFP expressing *Streptomyces* strain (Franco et al. 2017). To the best of our knowledge, our results show for the first time the presence of a streptomycete within an Arabidopsis root cell. Furthermore, the hyphae of coa1 appeared less electron dense in the endosphere than on the plant surface, which may reflect physiological differences between life inside and outside the plant. Still there are major gaps in our understanding of the endophytic biology of streptomycetes. For example, we did not find reproductive structures or spores within the endosphere. It therefore remains to be seen whether endophytes have a complete lifecycle inside plants or only remain in the vegetative growth phase, although we have to take into account that a seven day timeframe might not be sufficient to form spores within a plant. It has also been suggested that the endophytic lifestyle of actinobacteria may include the formation of small protoplast-like cells that lack a cell wall (Ramijan et al. 2016). These so-called L-forms may explain how the relatively large mycelial actinobacteria can still migrate and proliferate inside plant tissue. To further unravel the endophytic biology of *Streptomyces* we will focus future experiments on the identification of major genetic or morphological traits associated with the endophytic lifestyle.

### Response of endophytes to phytohormones

Actinobacteria play an important role in antibiosis and as probiotics to the plant due to the production of a diverse array of bioactive molecules (Viaene et al. 2016). We assessed the antimicrobial activity of the actinobacterial endophytes in overlay assays, using *Bacillus subtilis* (Gram-positive) and *Escherichia coli* (Gram-negative) as the target. By applying agar diffusion assays with the streptomycetes grown on minimal medium agar with mannitol and glycerol we showed that 11% of the isolates produced one or more compounds that inhibited growth of *E. coli* cells, while 14% inhibited growth of *B. subtilis*. Generally, the vast majority of actinomycetes isolated from soil samples are able to inhibit *B. subtilis* growth under routine laboratory conditions, which differs from what we find for our collection of endophytes. While the small numbers make it difficult to apply statistics, we cannot rule out that the reduced antibiotic activity against Gram-positive bacteria and in particular Firmicutes, may be typical of endophytic actinobacteria.

Genome sequencing revealed that actinobacteria have a lot of biosynthetic gene clusters for natural products that are poorly expressed (Kolter and van Wezel 2016; Nett et al. 2009). This offers a vast reservoir of potentially important bioactive molecules, including antibiotics. To allow screening of these compounds, strategies are required to elicit their expression (Rutledge and Challis 2015; van Wezel et al. 2009; Wu et al. 2015; Zhu et al. 2014a). We assessed the potential of phytohormones as elicitors, thereby mimicking the chemical environment of the rhizosphere and endosphere. For this, we exposed the endophytes to the plant hormones salicylic acid (SA), indole acetic acid (IAA, known as auxin) or jasmonic acid (JA). The strains were grown on minimal media, with or without 0.01%, 0.001% or 0.0001% (w/v) of either plant hormone. Square 12x12 cm agar plates were inoculated with spots from spore stocks of the endophytes. Interestingly, the percentage of strains exhibiting antibiotic activity roughly doubled, with 20% of the strains inhibiting *E. coli* cells, and 29% inhibiting growth of *B. subtilis* (Table S1). This effect could mostly be attributed to IAA, which had a more significant eliciting effect on the endophytes than SA or JA (Table S2). An example of elicited antimicrobial production by IAA is presented in Fig. S2. Additionally, we observed increased activity against the indicator strains as well, indicating either more production of the same antibiotic or production of a different (set of) antibiotic(s) (See Fig. S3). Although the concept of eliciting antimicrobial production by actinobacteria is already established, this experimental setup was connected to the biotic interactions between plant and streptomycete, aimed to mimic the chemical environment of the plant. Plant-endophyte interactions have likely played a key role in the evolution of the chemical diversity of actinomycete-derived natural products and signals that control the production of these antimicrobials are likely tied to the biotic interactions. This idea is further explored by the “cry for help” hypothesis, which additionally states that actinobacteria encounter trade-offs between the costs of producing complex natural products and their benefits, and may therefore produce these molecules specifically in response to ecological demands (van der Meij et al. 2017). To broaden our understanding of this concept it would be of interest to know which type(s) of compounds are produced in response to phytohormone exposure. Identification of the antimicrobials produced in response to phytohormones and metabolic networks will give better insight into the microbes’ responses to the plant’s “cry for help”.

## CONCLUSION

In this study, we show the recruitment of *Streptomyces* endophytes by *A. thaliana* Col-0 and *A. thaliana* Msl. Our pilot study shows that isolates falling within the groups of *S. olivochromogenes* and *S. clavifer* are overrepresented in the endosphere, suggesting that these endophytic streptomycetes may have specific characteristics that allow them to adapt to life inside the plant. This needs to be studied in more detail, and may also be extended to the study of other actinobacterial genera found in the endosphere. Additionally, we provide a first step towards the proof of concept for the ‘cry for help’ hypothesis, whereby plant hormones, in particular IAA, have a stimulating effect on antibiotic production by endophytic actinobacteria. These baseline experiments highlight the importance of exploring and exploiting plant-actinomycete interactions as elicitors for ‘antibiotic production on demand’.

## MATERIALS AND METHODS

### Bacteria, plants and growth conditions

*S. coelicolor* A3(2) M145 was obtained from the John Innes Centre strain collection in Norwich, UK, and *S. griseus* DSM40236 from the DSMZ culture collection (Braunschweig, Germany). *S. coelicolor* M145 and *S. griseus* DSM40236 were grown on SFM for 6 days at 30 °C, unless indicated differently. A sterile *A. thaliana* ecotype Columbia (Col-0) was used for the recruitment of endophytic actinobacteria under lab conditions. Plants were grown on a mixture of 9:1 substrate soil and sand (Holland Potgrond) at 21 °C, a 16h photoperiod, and 70% relative humidity. After 2 weeks of growth, the plants were harvested. Soil for the field experiment with *A. thaliana* ecotype Mossel (Msl) was collected in May 2016 at the Mossel area at the ‘Hoge Veluwe’ in the Netherlands (coordinates: N52°03’35.5” E5°45’06.4”), from four different spots within a radius of 100 m. If there was any vegetation present, the top 5–10 cm soil was removed. The soil was homogenised and all large parts such as dead roots and stones were removed. The soil was kept in a cold room at 4 °C until use. Seeds were sterilised in 4x diluted household bleach for 10 min, washed seven times with sterile MQ water, a short rinse with 70% ethanol and transferred to plates with a wet filter paper, placed at 4 °C for 48 hours and then moved to a 21 °C incubator in the dark. Mossel soil was placed in a tray with 3 x 3 cm pots and watered. To remove the endogenous seed population, the tray was placed in the greenhouse for 2 days. After weeding, the sterile seedlings on the plates were transplanted to the tray with Mossel soil and after 7 days the plants including the soil were planted into to the Mossel field. After 6 weeks of growth in the field, plants were harvested using a small shovel 3-4 cm around the base of the plant.

### Isolation of endophytes

*A. thaliana* Col-0 was surface sterilised by washing the plant 3 times in 70% EtOH, after which the plant material was crushed in MQ. Sterilization was confirmed by adding a sterilised plant onto LB agar, after which no bacterial growth was observed. Root tissue of *A. thaliana* Msl was cleaned, sonicated and ground with mortar and pestle in 1 mL phosphate buffer (per litre: 6.33 g of NaH_2_PO_4_·H_2_O, 10.96 g of Na_2_HPO_4_·H_2_O and 200 μL Silwet L-77). Both the crushed plant material and EC prep were spread onto the surface of a range of selective isolation media including humic acid agar (HA) (Hayakawa and Nonomura 1987), glucose casein agar (GCA) (Zhang 1985), soy flour mannitol medium (SFM) or minimal medium (MM) (Kieser et al. 2000). Initial selection was done on media supplemented with the antifungal agent nystatin (50 μg/ml) and the antibacterial agent nalidixic acid (10 μg/ml). Plates were incubated at 30°C for 4 to 25 days.

### Plant harvesting and DNA isolation of the microbial community

For *A. thaliana* Msl we applied the harvesting protocol as described before (Lundberg et al. 2012). In short, roots including rhizospheric soil were collected in a 50 mL tube containing 25 mL of phosphate buffer, and vortexed for 15 seconds. Replacing the buffer and vortexing was repeated until the buffer stayed clear. Roots were transferred to a 15 mL tube, sonicated (5 bursts of 30 seconds with 30 seconds breaks), vortexed, washed with phosphate buffer and dried on filter paper. Then, the roots were either ground for bacterial isolation, or flash frozen and stored at −80 °C for later DNA isolation. Four individual plants were pooled into one sample. DNA was isolated from soil and EC samples using the MoBio PowerSoil kit and the MP Bio Fast DNA spin kit, respectively. Quality and quantity of the DNA was checked by nanodrop and gel electrophoresis. Around 400 ng was sent for 16s rDNA sequencing at Beijing Genomics Institute (BGI).

### Amplicon sequencing

Using primers 515F (5’-GTGYCAGCMGCCGCGGTAA-3’) and 806R (5’-GGACTACNVGGGTWTCTAAT-3’), the V4 region was sequenced at BGI on the HiSeq2500 sequencing platform (Illumina). Raw data from BGI was processed using a previously reported custom implementation (Perez-Jaramillo et al. 2017) of QIIME (Caporaso et al. 2010) with minor modifications (described by Schneijderberg et al., in prep). In short, reads were quality filtered and filtered for chimeras using ChimeraSlayer. Using a 97% identity threshold, *de novo* OTUs were determined, which were taxonomically assigned using the RDP classifier 2.10 (Cole et al. 2014) with the GreenGenes database 28 (DeSantis et al. 2006). OTUs related to mitochondial and chloroplast sequences were removed, as were the OTUs that did not have 25 reads in at least 5 samples (“rare taxa”). To obtain relative abundance of the OTUs, the number of reads from a single OTU per sample was divided by the total number of reads of that sample after filtering for rare taxa.

### Analysis of actinobacteria based on 16S rRNA sequences

The 16S rRNA genes of the actinobacteria were amplified by PCR from liquid-grown mycelia using primers F1 (5’-GCGTGCTTAACACATGCAAG-3’) and R1 (5’-CGTATTACCGCGGCTGC T G-3’), which correspond to nt positions 15-34 and 465-483 of the 16S rRNA locus of *S. coelicolor* (van Wezel et al. 1991), respectively. PCRs were conducted as described (Colson et al. 2007) and sequenced using oligonucleotide F1. Sequencing was done at BaseClear in Leiden, the Netherlands. 16S rRNA gene analysis was performed using web based identify tool on EzBioCloud (https://www.ezbiocloud.net/identify). The identify service provides proven similarity-based searches against quality-controlled databases of 16S rRNA sequences. The top-hit information for each Identify Job was checked against *Streptomyces* focused MLSA based phylogenetic tree published elsewhere (Labeda et al. 2017).

### Microscopy

#### Light microscopy

Stereo microscopy was done using a Zeiss Lumar V12 microscope equipped with a AxioCam MRc, and confocal microscopy using a Zeiss Observer Microscope. For confocal microscopy, spores of the endophyte were added to the seeds. Seeds were put on ½ Murashige & Skoog medium with 1% sucrose and 0.8% agar and kept in the dark at 4 degrees for 48 hours. After the cold shock, seeds were incubated in the climate room for 1 week after which they were imaged. Samples were stained with propidium iodide 1:1000 (1 μg/ml). Samples were excited with laser light at a wavelength 535 nm to detect the propidium iodide.

#### Electron microscopy

Morphological studies on single colonies of endophytes by SEM were performed using a JEOL JSM6700F scanning electron microscope. For Streptomycete only samples, pieces of agar with biomass from 6-day-old colonies grown on SFM were cut and fixed with 1.5% glutaraldehyde (1 hour). Subsequently, samples were dehydrated (70% acetone 15 min, 80% acetone 15 min, 90% acetone 15 min, 100% acetone 15 min and critical point dried (Baltec CPD-030). Hereafter the samples were coated with gold using a gold sputter coater, and directly imaged using a JEOL JSM6700F. For SEM of Arabidopis samples, imaging timing was increased four-fold to optimise fixation and dehydration, and the 70% acetone step was done overnight.

Transmission electron microscopy (TEM) for the analysis of cross-sections of *Arabidopsis* roots and *Streptomyces* was performed with a JEOL1010 transmission electron microscope as described previously (Piette et al. 2005).

Samples were grown in the same way as for the light microscopy, and fixed with 1.5% glutaraldehyde for 4 hours. Post-fixation was performed with 1% Osmium tetroxide for 4 hours. Initial dehydration with 70% ethanol was done overnight. Hereafter using a high magnification (150x) stereo microscope (MZ16AF) root sections containing *Streptomyces* growth were selected, followed by dehydration in 1 hr steps (80%, 90%, 100% ethanol, 100% propylene oxide, 50/50 propylene oxide/EPON, 100% EPON). Subsequently samples were embedded in EPON and polymerised for 2 days at 60° Celsius. Ultrathin sections were cut on an ultramicrotome (Reichert Ultracut E), collected on copper grids and examined using a Jeol1010 transmission electron microscope at 70 kV.

### Antimicrobial activity assays

Antimicrobial activity of the endophytes was tested against *Bacillus subtilis* 168 or a derivative of Escherichia coli AS19-RrmA (Liu and Douthwaite 2002). Indicator bacteria were cultured in LB broth and incubated at 37 °C overnight. Antimicrobial assays were conducted using the double-layer agar method. Briefly, actinobacteria were inoculated on minimal medium agar plates containing mannitol and glycerol as the carbon sources (1% w/v), supplemented with either (+-)-jasmonic acid (Cayman chemical company, cas: 88-30-0), 3-indoleacetic acid (Sigma-Aldrich, cas: 87-51-4) or salycyclic acid (Alfa Aesar, cas: 69-72-7). The endophytes were typically incubated for 5 days at 30 °C, following which they were overlaid with LB soft agar (0.6% w/v agar) containing 300 μl of one of the indicator strains (OD 0.4 - 0.6), and then incubated overnight at 37 °C. The following day, antibacterial activity was determined by measuring the inhibition zones (mm) of the indicator strain surrounding the colonies.

## Acknowledgements

This work was supported by a grants 14218 and 14221 from the Netherlands Organization for Scientific research (NWO) to JR and GPvW, respectively. The authors declare no conflict of interests.

## REFERENCES

Anne J, Van Mellaert L, Eyssen H (1990) Optimum conditions for efficient transformation of *Streptomyces venezuelae* protoplasts. Appl Microbiol Biotechnol 32:431–435.

Barka EA, Vatsa P, Sanchez L, Gavaut-Vaillant N, Jacquard C, Klenk HP, Clément C, Oudouch Y, van Wezel GP (2016) Taxonomy, physiology, and natural products of the *Actinobacteria*. Microbiol Mol Biol Rev 80:1–43.

Bérdy J (2005) Bioactive microbial metabolites. J Antibiot (Tokyo) 58:1–26.

Bignell DRD, Seipke RF, Huguet-Tapia JC, Chambers AH, Parry RJ, Loria R (2010) *Streptomyces scabies* 87-22 Contains a Coronafacic Acid-Like Biosynthetic Cluster That Contributes to Plant-Microbe Interactions. Mol Plant-Microbe Interact 23:161–175.

Bonaldi M, Chen X, Kunova A, Pizzatti C, Saracchi M, Cortesi P (2015) Colonization of lettuce rhizosphere and roots by tagged *Streptomyces*. Front Microbiol 6:25.

Bukhalid RA, Chung SY, Loria R (1998) *nec1*, a gene conferring a necrogenic phenotype, is conserved in plant-pathogenic *Streptomyces* spp. and linked to a transposase pseudogene. Mol Plant-Microbe Interact 11:960–967.

Bulgarelli D, Rott M, Schlaeppi K, Ver Loren van Themaat E, Ahmadinejad N, Assenza F, Rauf P, Huettel B, Reinhardt R, Schmelzer E, Peplies J, Gloeckner FO, Amann R, Eickhorst T, Schulze-Lefert P (2012) Revealing structure and assembly cues for *Arabidopsis* root-inhabiting bacterial microbiota. Nature 488:91–95.

Caporaso JG, Kuczynski J, Stombaugh J, Bittinger K, Bushman FD, Costello EK, Fierer N, Pena AG, Goodrich JK, Gordon JI, Huttley GA, Kelley ST, Knights D, Koenig JE, Ley RE, Lozupone CA, McDonald D, Muegge BD, Pirrung M, Reeder J, Sevinsky JR, Turnbaugh PJ, Walters WA, Widmann J, Yatsunenko T, Zaneveld J, Knight R (2010) QIIME allows analysis of high-throughput community sequencing data. Nat Methods 7:335–336.

Cole JR, Wang Q, Fish JA, Chai B, McGarrell DM, Sun Y, Brown CT, Porras-Alfaro A, Kuske CR, Tiedje JM (2014) Ribosomal Database Project: data and tools for high throughput rRNA analysis. Nucleic Acids Res 42:D633–642.

Colson S, Stephan J, Hertrich T, Saito A, van Wezel GP, Titgemeyer F, Rigali S (2007) Conserved cis-acting elements upstream of genes composing the chitinolytic system of streptomycetes are DasR-responsive elements. J Mol Microbiol Biotechnol 12:60–66.

Doumbou CL, Akimov VV, Beaulieu C (1998) Selection and characterization of microorganisms utilizing thaxtomin A, a phytotoxin produced by *Streptomyces scabies*. Appl Environ Microbiol 64:4313–4316.

Edwards J, Johnson C, Santos-Medellin C, Lurie E, Podishetty NK, Bhatnagar S, Eisen JA, Sundaresan V (2015) Structure, variation, and assembly of the root-associated microbiomes of rice. Proc Natl Acad Sci U S A 112:E911–920.

Franco CM, Adetutu EM, Le HX, Ballard RA, Araujo R, Tobe SS, Paul B, Mallya S, Satyamoorthy K (2017) Complete Genome Sequences of the Endophytic *Streptomyces* sp. Strains LUP30 and LUP47B, Isolated from Lucerne Plants. Genome Announc 5.

Girard G, Traag BA, Sangal V, Mascini N, Hoskisson PA, Goodfellow M, van Wezel GP (2013) A novel taxonomic marker that discriminates between morphologically complex actinomycetes. Open Biol 3:130073.

Goodfellow M (2012) Phylum XXVI. Actinobacteria phyl. nov. In: Goodfellow M, Kämpfer P, Busse H-J et al. (eds) Bergey’s Manual of Systematic Bacteriology, vol 5. The Actinobacteria, Parts A and B, 2nd edn. Springer, New York, pp 1–2083. doi:10.1007/978-0-387-68233-4

Halim VA, Vess A, Scheel D, Rosahl S (2006) The role of salicylic acid and jasmonic acid in pathogen defence. Plant Biol 8:307–313.

Hayakawa M, Nonomura H (1987) Humic acid-vitamin agar, a new medium for selective isolation of soil actinomycetes. J Ferment Technol 65:501–509.

Hopwood DA (2007) *Streptomyces* in nature and medicine: the antibiotic makers. Oxford University Press, New York

Jourdan S, Francis IM, Kim MJ, Salazar JJ, Planckaert S, Frere JM, Matagne A, Kerff F, Devreese B, Loria R, Rigali S (2016) The CebE/MsiK Transporter is a Doorway to the Cello-oligosaccharide-mediated Induction of *Streptomyces scabies* Pathogenicity. Sci Rep 6:27144.

Keijser BJ, van Wezel GP, Canters GW, Kieser T, Vijgenboom E (2000) The ram-dependence of *Streptomyces lividans* differentiation is bypassed by copper. J Mol Microbiol Biotechnol 2:565–574.

Kieser T, Bibb MJ, Buttner MJ, Chater KF, Hopwood DA (2000) Practical *Streptomyces* Genetics Norwich, UK John Innes Foundation.

Kim TU, Cho SH, Han JH, Shin YM, Lee HB, Kim SB (2012) Diversity and physiological properties of root endophytic actinobacteria in native herbaceous plants of Korea. J Microbiol 50:50–57.

Kolter R, van Wezel GP (2016) Goodbye to brute force in antibiotic discovery? Nat Microbiol 1:15020.

Kortemaa H, Rita H, Haahtela K, Smolander A (1994) Root Colonization Ability of Antagonistic *Streptomyces griseoviridis*. Plant Soil 163:77–83.

Kruasuwan W, Salih TS, Brozio S, Hoskisson PA, Thamchaipenet A (2017) Draft Genome Sequence of Plant Growth-Promoting Endophytic *Streptomyces* sp. GKU 895 Isolated from the Roots of Sugarcane. Genome Announc 5.

Labeda DP, Dunlap CA, Rong X, Huang Y, Doroghazi JR, Ju KS, Metcalf WW (2017) Phylogenetic relationships in the family Streptomycetaceae using multi-locus sequence analysis. Antonie Van Leeuwenhoek 110:563–583.

Lambert S, Traxler MF, Craig M, Maciejewska M, Ongena M, van Wezel GP, Kolter R, Rigali S (2014) Altered desferrioxamine-mediated iron utilization is a common trait of bald mutants of *Streptomyces coelicolor*. Metallomics 6:1390–1399.

Lebeis SL, Paredes SH, Lundberg DS, Breakfield N, Gehring J, McDonald M, Malfatti S, Glavina del Rio T, Jones CD, Tringe SG, Dangl JL (2015) Salicylic acid modulates colonization of the root microbiome by specific bacterial taxa. Science 349:860–864.

Lee SO, Choi GJ, Choi YH, Jang KS, Park DJ, Kim CJ, Kim JC (2008) Isolation and characterization of endophytic actinomycetes from Chinese cabbage roots as antagonists to Plasmodiophora brassicae. J Microbiol Biotechnol 18:1741–1746.

Liu M, Douthwaite S (2002) Activity of the ketolide telithromycin is refractory to Erm monomethylation of bacterial rRNA. Antimicrob Agents Chemother 46:1629–1633.

Loria R, Kers J, Joshi M (2006) Evolution of plant pathogenicity in *Streptomyces*. Annu Rev Phytopathol 44:469–487.

Lundberg DS, Lebeis SL, Paredes SH, Yourstone S, Gehring J, Malfatti S, Tremblay J, Engelbrektson A, Kunin V, del Rio TG, Edgar RC, Eickhorst T, Ley RE, Hugenholtz P, Tringe SG, Dangl JL (2012) Defining the core *Arabidopsis thaliana* root microbiome. Nature 488:86–90.

Misk A, Franco C (2011) Biocontrol of chickpea root rot using endophytic actinobacteria. BioControl:811–822.

Nett M, Ikeda H, Moore BS (2009) Genomic basis for natural product biosynthetic diversity in the actinomycetes. Nat Prod Rep 26:1362–1384.

Perez-Jaramillo JE, Carrion VJ, Bosse M, Ferrao LFV, de Hollander M, Garcia AAF, Ramirez CA, Mendes R, Raaijmakers JM (2017) Linking rhizosphere microbiome composition of wild and domesticated *Phaseolus vulgaris* to genotypic and root phenotypic traits. ISME J 11:2244–2257.

Piette A, Derouaux A, Gerkens P, Noens EE, Mazzucchelli G, Vion S, Koerten HK, Titgemeyer F, De Pauw E, Leprince P, van Wezel GP, Galleni M, Rigali S (2005) From dormant to germinating spores of *Streptomyces coelicolor* A3(2): new perspectives from the *crp* null mutant. J Proteome Res 4:1699–1708.

Ramijan K, Willemse J, Ultee E, Wondergem J, van der Meij A, Briegel A, Heinrich A, van Wezel GP, Claessen D (2016) Reversible metamorphosis in a bacterium. BioRXiv.

Rivas-San Vicente M, Plasencia J (2011) Salicylic acid beyond defence: its role in plant growth and development. J Exp Bot 62:3321–3338.

Rutledge PJ, Challis GL (2015) Discovery of microbial natural products by activation of silent biosynthetic gene clusters. Nat Rev Microbiol 13:509–523.

Seipke RF, Kaltenpoth M, Hutchings MI (2012) *Streptomyces* as symbionts: an emerging and widespread theme? FEMS Microbiol Rev 36:862–876.

Singh SP, Gaur R (2016) Evaluation of antagonistic and plant growth promoting activities of chitinolytic endophytic actinomycetes associated with medicinal plants against *Sclerotium rolfsii* in chickpea. J Appl Microbiol 121:506–518.

van de Mortel JE, de Vos RCH, Dekkers E, Pineda A, Guillod L, Bouwmeester K, van Loon JJA, Dicke M, Raaijmakers JM (2012) Metabolic and Transcriptomic Changes Induced in *Arabidopsis* by the Rhizobacterium *Pseudomonas fluorescens* SS101. Plant Physiology 160:2173–2188.

van der Meij A, Worsley SF, Hutchings MI, van Wezel GP (2017) Chemical ecology of antibiotic production by actinomycetes. FEMS Microbiol Rev 41:392–416.

van Dissel D, Claessen D, Van Wezel GP (2014) Morphogenesis of *Streptomyces* in submerged cultures. Adv Appl Microbiol 89:1–45.

van Wezel GP, McKenzie NL, Nodwell JR (2009) Chapter 5. Applying the genetics of secondary metabolism in model actinomycetes to the discovery of new antibiotics. Methods Enzymol 458:117–141.

van Wezel GP, Vijgenboom E, Bosch L (1991) A comparative study of the ribosomal RNA operons of Streptomyces coelicolor A3(2) and sequence analysis of rrnA. Nucleic Acids Res 19:4399–4403.

Viaene T, Langendries S, Beirinckx S, Maes M, Goormachtig S (2016) *Streptomyces* as a plant’s best friend? FEMS Microbiol Ecol 92.

Wu C, Kim HK, van Wezel GP, Choi YH (2015) Metabolomics in the natural products field - a gateway to novel antibiotics. Drug Discov Today Technol 13:11–17.

Zhang J (1985) Microbial Taxonomy. Fudan University Press:214–218.

Zhao Y (2010) Auxin biosynthesis and its role in plant development. Annu Rev Plant Biol 61:49–64.

Zhu H, Sandiford SK, van Wezel GP (2014a) Triggers and cues that activate antibiotic production by actinomycetes. J Ind Microbiol Biotechnol 41:371–386.

Zhu H, Swierstra J, Wu C, Girard G, Choi YH, van Wamel W, Sandiford SK, van Wezel GP (2014b) Eliciting antibiotics active against the ESKAPE pathogens in a collection of actinomycetes isolated from mountain soils. Microbiology 160:1714–1725.

